# Endothelial Piezo1 regulates muscle AKT abundance and mechanically induced muscle hypertrophy

**DOI:** 10.1101/2025.09.02.673617

**Authors:** Fiona Bartoli, Peter Tickle, Harrison Gallagher, Marjolaine Debant, Chew W Cheng, T Simon Futers, Helene Daou, Nadira Y Yuldasheva, David J Beech, T Scott Bowen

## Abstract

Piezo1 proteins form mechanically activated nonselective cation channels with key roles in endothelial responses to mechanical forces. Here we reveal that endothelial Piezo1 regulates skeletal muscle hypertrophy following mechanical overload. Using a conditional endothelial cell-specific deletion of Piezo1 in adult mice, we assessed the role of endothelial Piezo1 in mechanical overload-induced muscle hypertrophy in the extensor digitorum longus. Endothelial Piezo1 deletion blunted muscle hypertrophy following mechanical overload despite normal baseline vascularisation, as evidenced by a lack of increase in muscle mass and abolished myofibre growth. Despite being required for optimal muscle growth, endothelial Piezo1 was dispensable for muscle regeneration after injury. The reduced muscle growth following mechanical overload was not associated with impaired myonuclear accretion but rather with reduced abundance of AKT (protein kinase B), a key regulator of muscle protein synthesis. We suggest that endothelial Piezo1 participates in a myofibre partnership that regulates skeletal muscle growth in response to mechanical overload.

**Significance:** What allows skeletal muscle to grow in response to increased physical demand is relatively poorly understood. The work in this paper suggests that a mechanically activated ion channel in vascular endothelial cells, termed Piezo1, is required for muscle growth in response to increased mechanical load, regulating a major pathway of myofiber hypertrophy. A new concept is thus proposed in which a vascular mechanical axis is a key determinant of muscle growth, ensuring muscle adaptation matches cardiovascular capacity. This new understanding may inform new ways to address health problems associated with sedentary lifestyles and ageing.

## INTRODUCTION

Skeletal muscle, the largest tissue mass in the human body, contains multiple cell subtypes (including myofibres, endothelial, pericyte, immune, and muscle stem cells [MuSC]) that integrate external cues vital for supporting movement, metabolism, and overall quality of life ^1^. In health, increased mechanical loading (e.g., resistance exercise) triggers biochemical signalling to increase muscle growth (hypertrophy)^2^, whereas ageing and disease are characterised by muscle atrophy^1^. Although regular exercise improves skeletal muscle health and quality of life, the fundamental mechanisms directing these benefits are only partly understood.

Increased mechanical loading on skeletal muscle triggers biochemical signalling that increases muscle growth^3^. Muscle growth due to resistance-dependent exercise is thought to be determined primarily by: MuSC division and fusion to add new myonuclei to existing myofibres (myonuclear accretion); and myofibre protein synthesis^2^. Many upstream and distinct mechanisms regulate muscle growth including hormonal/growth factors (e.g., IGF-1, β2-agonists) and metabolic cues^3^. Resistance exercise induces muscle hypertrophy in a load-dependent manner^2^, which also indicates a coupling between mechanical load and myofibre growth. However, the mechanosensing mechanisms involved in myofibre growth under tension remain mysterious^4^ and only a limited number of candidate mediators have been identified. These include myofibre plasma membrane transduction, such as G protein coupled receptor 56 signalling and its extracellular ligand collagen type III (Col3a1)^5^, diacylglycerol kinase-ζ activation^6^, JNK/SMAD axis^7^, and nitric oxide (NO)-Ca^2+^ signalling^8^; but also TRIM28 expression^9^ and primary cilia sensing in MuSCs^10^. An important mechanosensing mechanism could also reside within the local muscle vascular niche, via endothelial cells, by direct crosstalk with myofibres and MuSCs to regulate subsequent muscle growth^3^. Muscle mass is closely correlated to blood flow^11^ and emerging evidence supports the idea that endothelial cells may coordinate muscle mass regulation^12^, however whether this acts via mechanosensory mechanisms is unclear.

Piezo1 is a calcium-permeable channel subunit widely expressed in the body with a crucial role in sensing mechanical stimuli and regulating cell behaviours, such as proliferation, migration, and apoptosis^13^. It is vital for vasculature development, lymphatic development and blood pressure regulation^14–19^. The deletion of Piezo1 in various cell types from the musculoskeletal system is linked to bone health^20,21^ and tendon stiffness^22,23^. Piezo1 expression in myofibers^24^ and MuSCs^13,25^ influences skeletal muscle atrophy and regeneration during immobilisation or injury. However, little is known about whether Piezo1 controls muscle growth in response to mechanical loads. Despite a ubiquitous expression, a major site of expression of Piezo1 is the endothelium^26^, which consists of a monolayer of endothelial cells lining the inner surface of blood vessels and lymphatics. Adult mice deficient for Piezo1 specifically in the endothelium displayed reduced muscle capillarity alongside lower physical running performance^27^.

Here, we investigated the role endothelial Piezo1 plays in regulating adult muscle growth using a mechanical overload-dependent model of skeletal muscle hypertrophy^3^. We suggest that endothelial Piezo1 is required for optimal muscle growth following mechanical overload independent of capillary density. Overall, our data suggest that endothelial Piezo1 is a pivotal mechanosensitive mechanism of muscle hypertrophy, which could offer a potential therapeutic target for increasing muscle mass in relevant populations such as in ageing, disease, immobilisation, or with natural genetic variations.

## MATERIALS AND METHODS

### Study approval

All animal use was authorized by the University of Leeds Animal Ethics Committee and The Home Office, UK.

### Piezo1^ΔEC^ mouse model

*Piezo1* conditional KO mice were generated at the University of Leeds by crossing C57BL/6J Piezo1 floxed mice (*Piezo1^flox/flox^*) with a Cre transgenic line driven by the Cadherin-5 promoter (Tg(Cdh5-Cre/ERT2)1Rha) to generate Cre-positive, loxP-homozygous (*Piezo1^flox/flox^*/Cdh5-Cre) conditional KO mice as described previously^17,19,27^. The disruption of *Piezo1* gene specifically in the endothelium of *Piezo1^flox/flox^*/Cdh5-Cre mice was induced at age 8-10 weeks by consecutive tamoxifen injections (75 mg.kg^−1^ for 5 days; IP). *Piezo1* conditional KO mice are called *Piezo1^ΔEC^* mice. Cre-negative littermates received tamoxifen injections and are referred as to control mice. Housing and genotyping of the mice were carried out as previously described^27^. Male mice were used for experiments after either 2 weeks or 10 weeks deletion period of *Piezo1* gene as previously described^17,27^.

### Overload model of muscle hypertrophy

*Piezo1^ΔEC^* mice and their control littermates were submitted to unilateral mechanical overload of the extensor digitorum longus (EDL) muscle by removal of the synergist muscle tibialis anterior (TA) as previously described^28^. Anaesthesia was induced and maintained with isoflurane (5% and 2% respectively, in 100% O_2_). The distal tendon of the TA muscle was sectioned, then the TA was carefully dissected and excised. The contralateral limb served as an internal control. Removal of the TA muscle increased the load burden on the EDL muscle, causing an overload-induced hypertrophy of the muscle. Mice were sacrificed 14 days after the overload surgery.

### Model of muscle regeneration

*Piezo1^ΔEC^* mice and control littermates were submitted to muscle injury while anaesthetised with isoflurane as for the overload surgery. Mice received an intramuscular injection of 1.2% barium chloride (BaCl_2_, in 50 μL saline) into the right TA muscle. The left TA muscle was injected with saline solution to serve as an internal control. Mice were sacrificed 7 days after injection.

### Sampling

Mice were sacrificed under terminal anaesthesia in accordance with the Schedule 1 Code of Practice, UK Animals Scientific Procedures Act 1986. Skeletal muscles of interest were dissected from both hindlimbs, cleaned, dried, weighed, and divided into two parts cross-sectionally. The distal part was embedded in Tissue-Tek O.C.T. compound and frozen in isopentane cooled by liquid nitrogen for histology and immunohistochemistry studies, as previously described^28^. The proximal part was snap frozen directly in liquid nitrogen for molecular biology studies. Tissues were stored at −80°C until analysis.

### Histological and immunohistochemical analysis

For overload model, serial transverse cryosections of 10 μm thickness of EDL were cut using a cryostat set at −20°C (Leica) and mounted on SuperFrost Plus Adhesion slides (ThermoFisher Scientific, 10149870) to determine fibre size, capillarity, nuclei density and fibre types. For fibre type, immunostainings were performed as previously described^27^. After blocking with 10% goat serum in PBS, sections were incubated 2h at room temperature with the following primary antibody cocktails: (a) mouse IgG2b monoclonal anti–MHC type 1 (BA-F8), mouse IgG1 monoclonal anti–MHC type 2a (SC-71), and mouse IgM monoclonal anti–MHC type 2b (BF-F3); or (b) mouse IgG1 monoclonal anti–MHC type 2a (SC-71) and mouse IgM monoclonal anti–MHC type 2x (6H1). All antibodies (DSHB, University of Iowa) were used at 0.5 μg/mL. Secondary antibody cocktails were used as follow: (a) DyLight 405 IgG2b goat anti-mouse (Jackson Immunoresearch, 115-475-207; 1:500), Alexa Fluor 488 IgG1 goat anti-mouse (ThermoFisher Scientific, A-21121; 1:500), and Alexa Fluor 555 IgM goat anti-mouse (ThermoFisher Scientific, A-21426; 1:500); or (b) Alexa Fluor 488 IgG1 goat anti-mouse (1:500) and Alexa Fluor 555 IgM goat anti-mouse (1:500). Sections were cover-slipped using ProLong Gold Antifade Mountant (ThermoFisher Scientific, P36934).

For fibre size, capillarity and myonuclei density, immunostainings were performed as previously described^27^. Sections were fixed for 10min with 4% paraformaldehyde (PFA), permeabilized for 10min with 0.1% Triton X-100, blocked for 1h with 5% goat serum and 1% BSA, then incubated overnight at 4°C with rat anti-CD31 (BD Biosciences, 550274; 1:100) to detect blood vessels. The secondary antibody cocktails used was Alexa Fluor 647 chicken anti-rat (ThermoFisher Scientific, A-21472; 1:500) in combination with CF^®^488A-conjugated wheat germ agglutinin (WGA) (Biotium, Fremont, CA, USA, 29022-1, 1:100) to label cell membranes for 1h at room temperature. Sections were coverslipped using ProLong Gold Antifade Mountant with DAPI (ThermoFisher Scientific, P36935) to counterstain nuclei. For regeneration model, muscle histology was assessed on serial transverse cryosections (10 μm) of TA, stained with hematoxylin and eosin. Slides were imaged at 10x magnification using the Zeiss Axioscan Slide Scanner (Thornwood, NY). Two regions of interest (∼100 fibres each) were taken for each image. Regenerating fibres with centralised nuclei were quantified manually using ImageJ software.

### Statistics

The number (N) of mice studied per experiment is indicated in figure legends. All data are presented as mean ± SD. Outliers were tested and excluded if necessary, according to the results of ROUT test (Q = 1%) performed in GraphPad Prism software (v10, San Diego, CA, USA). Statistical significance was evaluated with unpaired Student’s t test when comparing 2 groups, and with 2-way ANOVA followed by post hoc Tukey’s test or Šídák correction for multiple comparisons, as stated in figure legends. For all experiments, a P value of less than 0.05 was considered significant.

## RESULTS

### Deletion of endothelial Piezo1 blunts load-induced muscle growth

To test whether endothelial Piezo1 supports load-induced muscle growth, endothelial-specific disruption of Piezo1 was induced in adult mice at 8 weeks of age (*Piezo1^ΔEC^* mice)^17,27^. A deletion period of 2 weeks was used before *Piezo1^ΔEC^* mice and control littermates (Ctrl mice) were exposed to mechanical overload to induce muscle growth^3^. The model included unilateral removal of the tibialis anterior (TA) muscle to induce hypertrophy of the synergist extensor digitorum longus (EDL)^28^. The EDL from hypertrophied leg and contralateral leg (internal control) were harvested 2 weeks after mechanical overload and thus when the mice were 12 weeks old. No differences in body weight (Figure 1A) or tibia length (Figure 1B) were found in *Piezo1^ΔEC^* mice at 12 weeks compared to Ctrl mice. In contrast, the increase in EDL wet-mass after overload surgery was reduced in *Piezo1^ΔEC^* mice (Figure 1C-D). This reduction was still observed when EDL wet-mass was corrected for body mass or tibia length (Figure 1E-F). Following overload, EDL wet mass increased by 35% in Ctrl mice but only 20% in *Piezo1^ΔEC^* mice (Figure 1G). Results were similar after a longer Piezo1 deletion period of 10 weeks and analysis of the mice at 22 weeks of age (Supplementary Figure 1). The data suggest that loss of endothelial Piezo1 impairs the ability of skeletal muscle to increase mass in response to mechanical loading.

**Figure 1:**
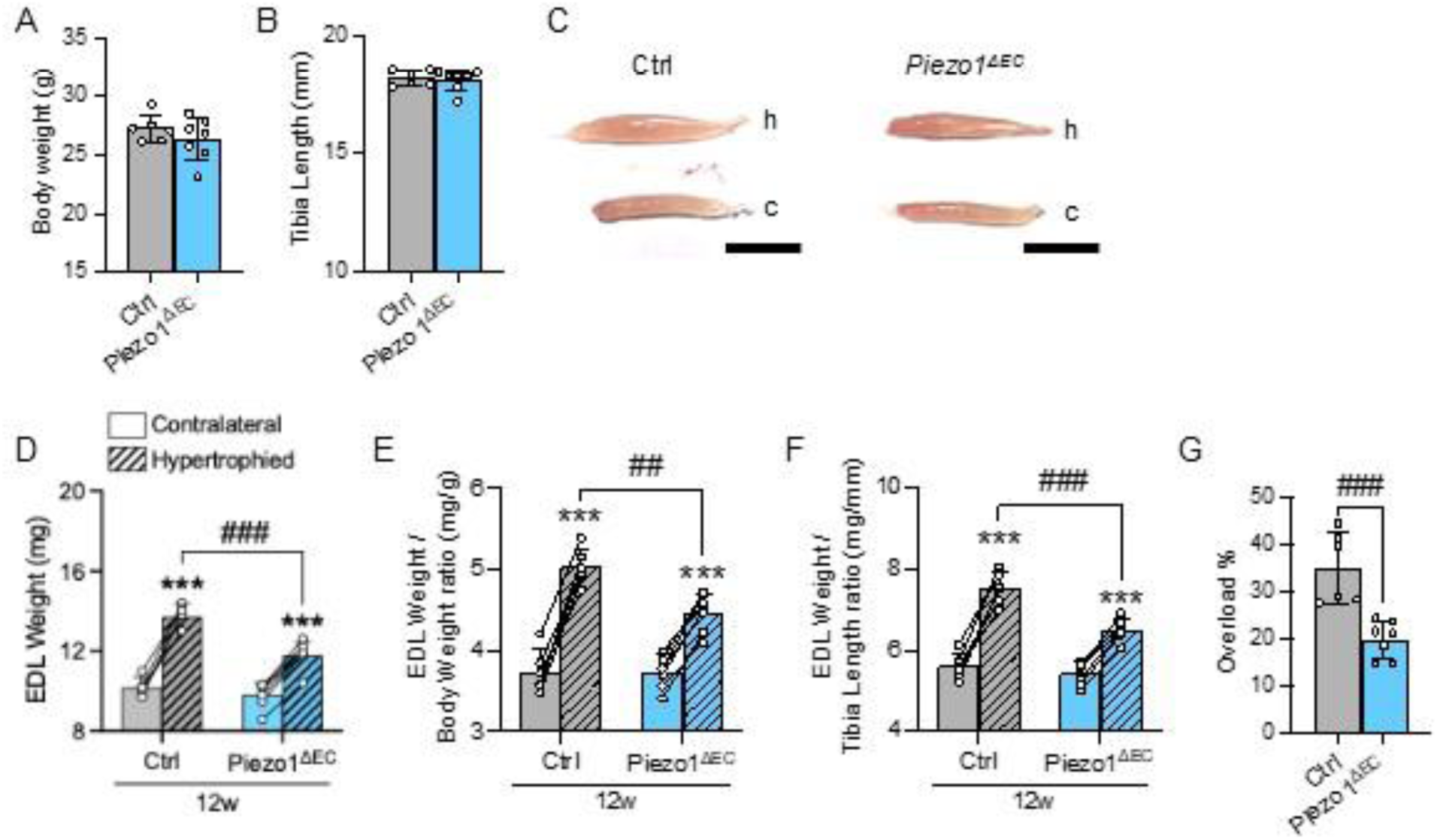
Two-week deletion of endothelial Piezo1 decreases muscle growth capacity. Data in grey are for 12-weeks old Ctrl mice and data in blue are for matched *Piezo1^ΔEC^* mice. Full bars are for contralateral leg (c, control) and hatched bars are for hypertrophied leg (h, overload surgery). A, Body weight; B, Tibia length; C, Representative morphology images of dissected EDL muscles (scale bar 0.5 cm); D, EDL wet-muscle mass; E, EDL mass to body weight ratio; F, EDL weight to tibia length ratio; G, Relative increase in muscle mass following overload compared to contralateral leg. Data are for N = 6-7 mice per group (mean ± S.D.). Superimposed white circles are the individual underlying data values and lines are direct comparison between contralateral and hypertrophied legs for each individual mouse. \*\*\**P*<0.001 vs. contralateral leg, ^##^*P*<0.01, ^###^*P*<0.001 vs. ctrl mice. Statistical significance was evaluated using Student’s *t* test (A-B, G) or 2-way ANOVA followed by Tukey’s HSD post hoc test for multiple comparisons (D-F).

### Load-induced myofibre hypertrophy is abolished after endothelial Piezo1 deletion

Muscle wet mass provides only a gross index of muscle hypertrophy limiting mechanistic insights^3^. Therefore, we next directly assessed changes related to myofibre phenotype in terms of size (cross-sectional area; CSA), density, and isoforms using stained cryosections in Ctrl and *Piezo1^ΔEC^* mice following mechanical overload. While the average myofiber CSA increased in Ctrl mice after mechanical overload, myofiber hypertrophy was absent in EDL muscle from *Piezo1^ΔEC^* mice (Figure 2A-B). Ctrl mice showed a decreased fibre density after overload, consistent with the increased myofibre CSA, which was not found in *Piezo1^ΔEC^* mice (Figure 2C). To further confirm the presence of cellular hypertrophy after mechanical overload, we next examined the fibre size distribution. Ctrl mice showed a decreased fibre <1000 μm^2^ frequency and an increased >1000 μm^2^ frequency after overload, whereas *Piezo1^ΔEC^* mice exhibited no change in either parameter (Figure 2D-F). Myofibre hypertrophy can also be influenced by a fibre-type shift. There were no shifts in fibre types in either group (Figure 3A-B) but type 2X and 2B fibre hypertrophy in the Ctrl EDL was not seen in the *Piezo1^ΔEC^* EDL (Figure 3C).

**Figure 2:**
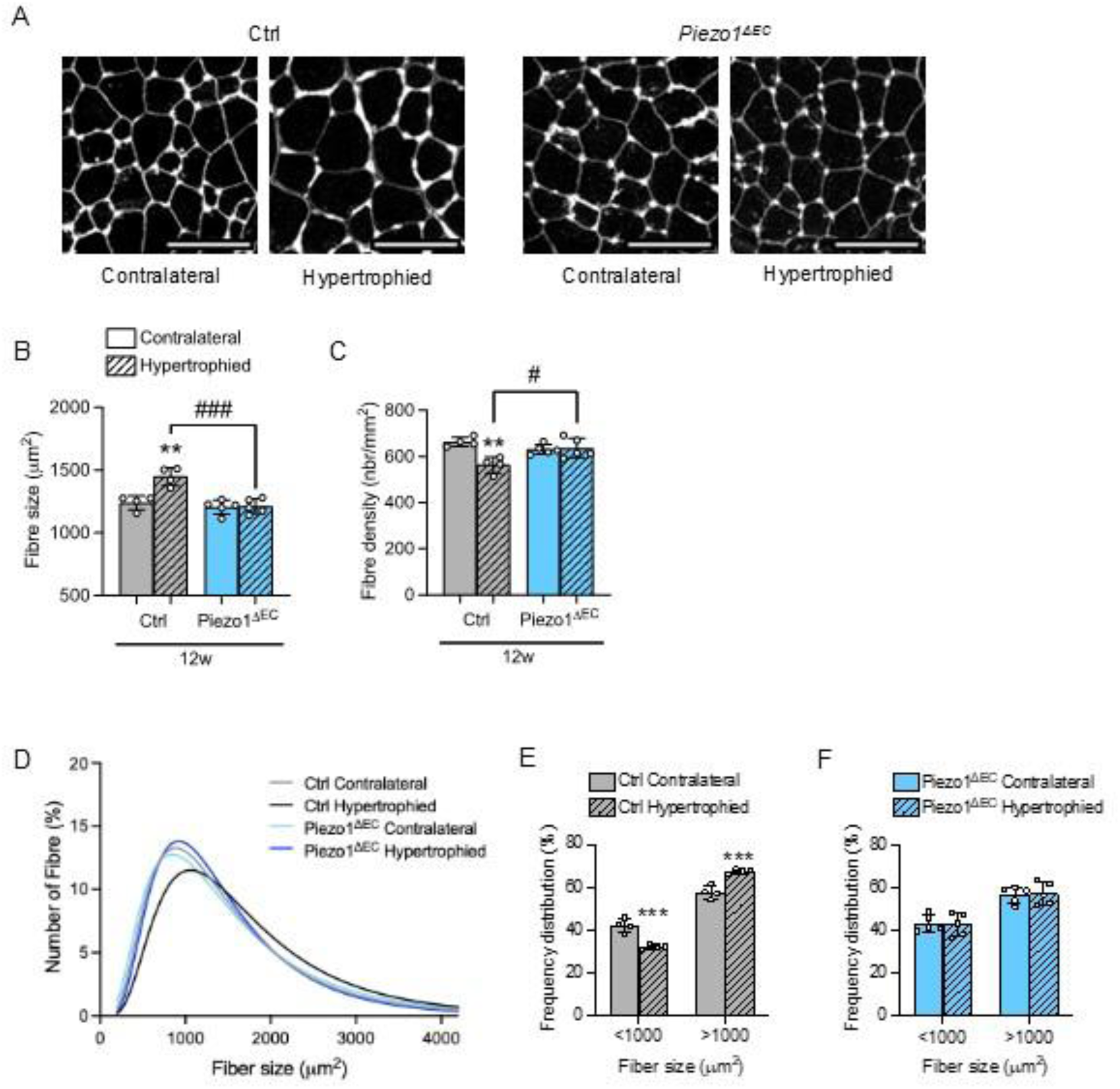
Two-week deletion of endothelial Piezo1 blunts overload-induced myofibre hypertrophy. Data in grey are for 12-weeks old Ctrl mice and data in blue are for matched *Piezo1^ΔEC^*mice. Full bars are for contralateral leg (c, control) and hatched bars are for hypertrophied leg (h, overload surgery). A, Immunohistochemistry for staining by wheat germ agglutinin (WGA, white) in contralateral and hypertrophied EDL muscle sections of Ctrl and *Piezo1^ΔEC^* mice. Scale bars, 100 μm. B, Fibre size determination using the mean cross-sectional area (CSA); C, Fibre density; D, Fibre size distribution, E-F, Mean frequency of fibre distribution for Ctrl mice (E) and *Piezo1^ΔEC^* mice (F). Data are for N = 4-5 mice per group (mean ± S.D.). Superimposed dots are the individual underlying data values for each individual mouse. \*\**P*<0.01, \*\*\**P*<0.001 vs. contralateral leg, ^#^*P*<0.05, ^###^*P*<0.001 vs. ctrl mice. Statistical significance was evaluated using 2-way ANOVA followed by Tukey’s HSD post hoc test (B-C) or Šídák correction (E-F) for multiple comparisons.

**Figure 3:**
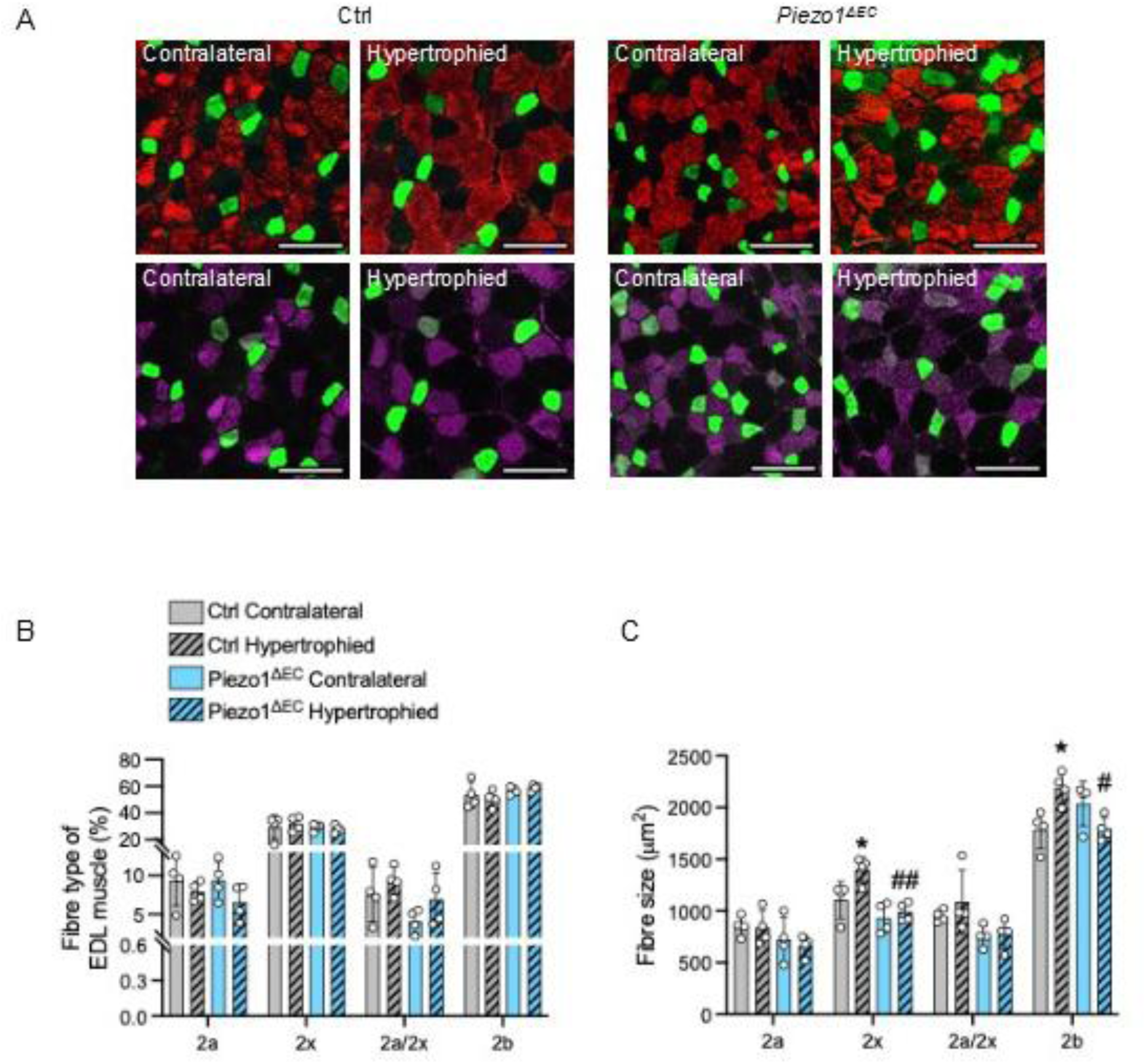
Two-week deletion of endothelial Piezo1 does not change fibre types. Data in grey are for 12-weeks old Ctrl mice and data in blue are for matched *Piezo1^ΔEC^* mice. Full bars are for contralateral leg (c, control) and hatched bars are for hypertrophied leg (h, overload surgery). A, Immunohistochemistry of EDL muscle sections of Ctrl and *Piezo1^ΔEC^* mice for myosin heavy chain (MHC) type 2a (green) plus type 2b (red, top) or type 2x (magenta, bottom). Scale bars, 100 μm. B, Quantification of the relative frequency of the different fibre types in EDL muscles. C, Fibre size determination using the mean cross-sectional area of the different fibre types in EDL muscles. Data are for N = 4 mice per group (mean ± S.D.). Superimposed dots are the individual underlying data values for each individual mouse. \**P*<0.05 vs. contralateral leg, ^#^*P*<0.05, ^##^*P*<0.01 vs. ctrl mice. Statistical significance was evaluated using 2-way ANOVA followed by Tukey’s HSD post hoc test for multiple comparisons

Given 10-week endothelial Piezo1 deletion was previously found to impair baseline muscle capillarity^27^ and this could potentially limit load-induced muscle growth^3,29^, we next examined changes in EDL capillary density and capillary-to-fibre (C:F) ratio 2 weeks following Piezo1 deletion. At this 2-week time-point, the capillary density was not different between Ctrl and *Piezo1^ΔEC^* EDL (Figure 4A-C), and while basal C:F ratio was not different between groups, overload increased the C:F ratio in Ctrl mice by 20% compared to the smaller and non-significant 9% increase found in *Piezo1^ΔEC^*mice (Figure 4C), supporting past reports that muscle growth/mass is closely coupled to vascular adaptations^29–31^. Taken together, these data indicate endothelial

**Figure 4:**
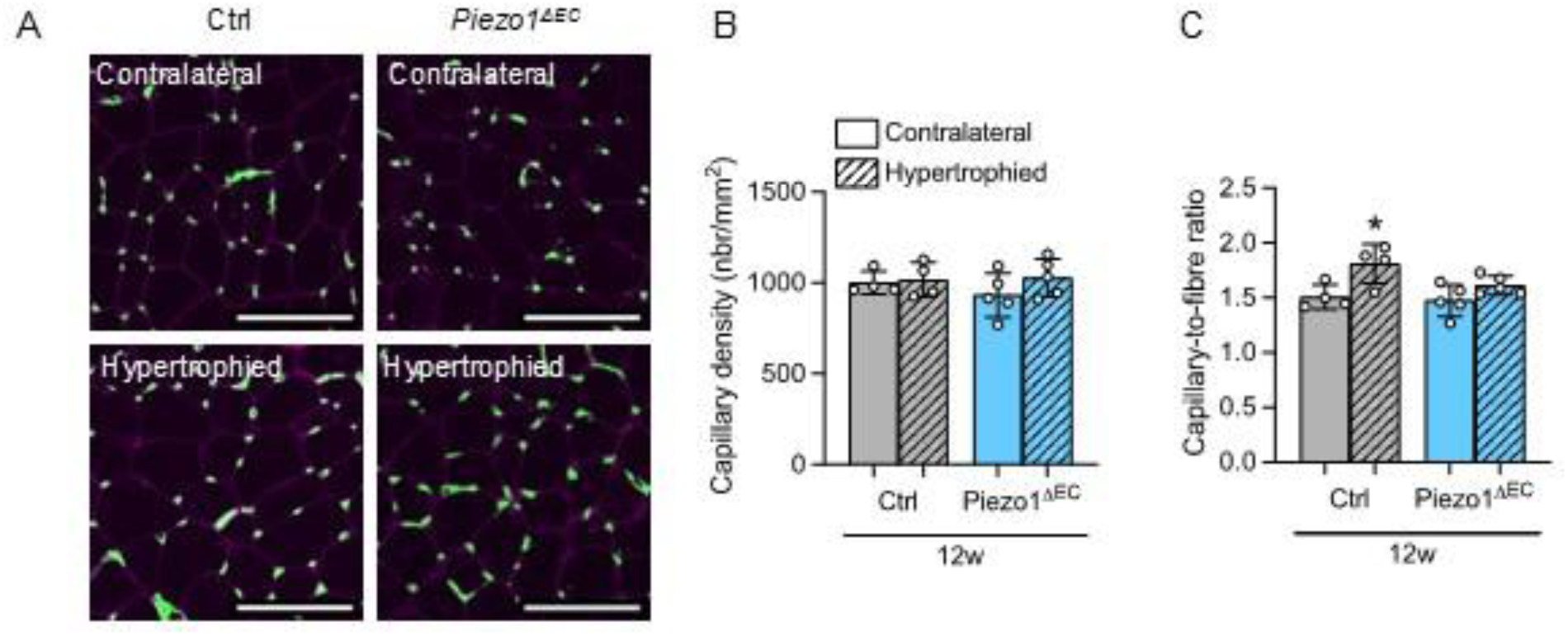
Two-week deletion of endothelial Piezo1 does not change capillary density. Data in grey are for 12-weeks old Ctrl mice and data in blue are for matched *Piezo1^ΔEC^* mice. Full bars are for contralateral leg (c, control) and hatched bars are for hypertrophied leg (h, overload surgery). A, Immunohistochemistry of EDL muscle sections of Ctrl and *Piezo1^ΔEC^*mice for capillaries (CD31, green). Scale bars, 100 μm. B, Capillary density; C, Capillary-to-fibre ratio. Data are for N = 4-5 mice per group (mean ± S.D.). Superimposed dots are the individual underlying data values for each individual mouse. \**P*<0.05 vs. contralateral leg. Statistical significance was evaluated using 2-way ANOVA followed by Tukey’s HSD post hoc test for multiple comparisons.

Piezo1 deletion impairs load-induced muscle mass by abolishing myofibre growth response in Type 2X and 2B fibres independent of baseline muscle capillarity.

### Endothelial Piezo1 deletion does not impair myonuclear accretion or muscle regeneration after injury

Muscle fibre hypertrophy can depend on MuSC-dependent myonuclear accretion^32–34^. We therefore evaluated myonuclear accretion in Ctrl and *Piezo1^ΔEC^* EDL following mechanical overload. Both groups similarly increased myonuclear density and nuclei-to-fibre ratio (Figure 5A-C). To further investigate myonuclear accretion, we used an injury model^35^ that require MuSCs to facilitate muscle regeneration^36^. Ctrl and *Piezo1^ΔEC^* TA muscles were submitted to barium chloride (BaCl_2_)-induced injury and resulting fibre necrosis, which activates MuSCs. Seven days post-injection, TA weight was lower in BaCl_2_-injected leg compared to vehicle-injected leg but there were no differences between Ctrl and *Piezo1^ΔEC^* mice (Figure 6A-B, Table 1). In addition, CSA of regenerated fibres was similar between genotypes (Figure 6C-D), suggesting endothelial Piezo1 was not supporting the regenerative process after muscle injury.

**Figure 5:**
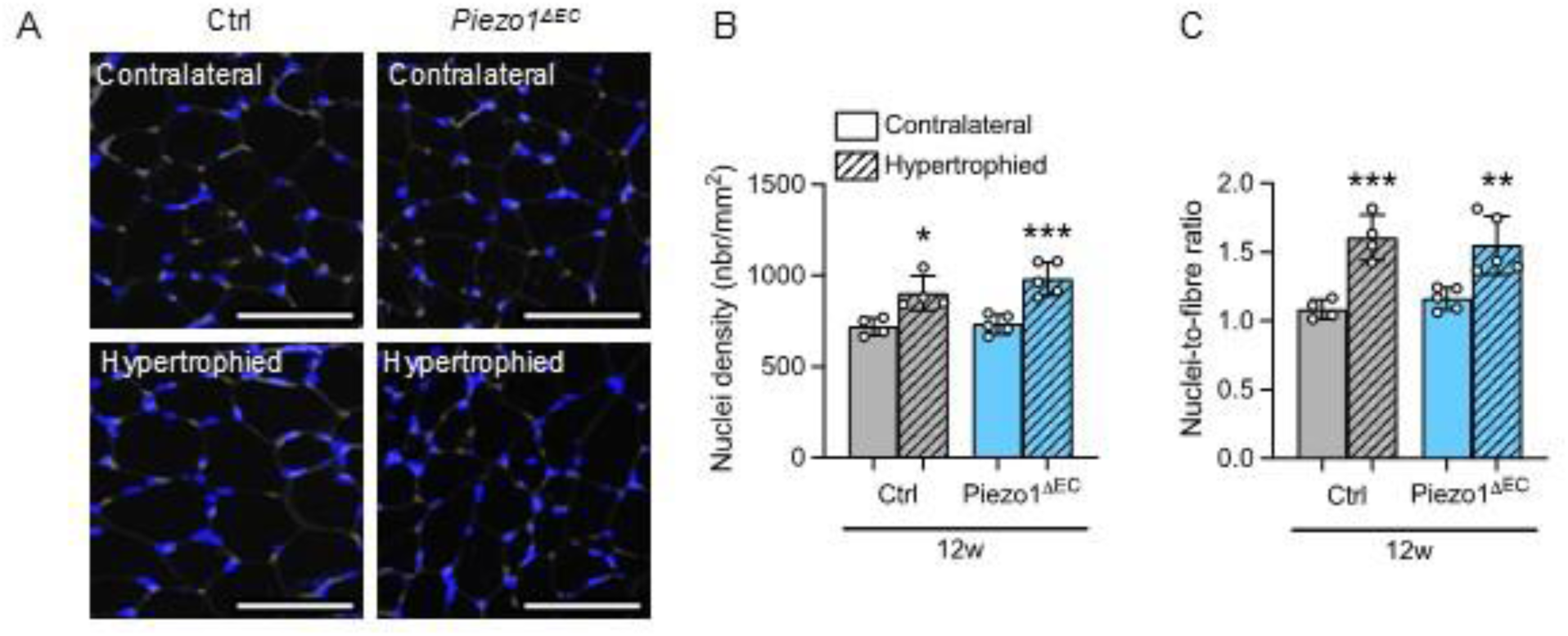
Two-week deletion of endothelial Piezo1 does not change overload-induced myonuclear accretion. Data in grey are for 12-weeks old Ctrl mice and data in blue are for matched *Piezo1^ΔEC^* mice. Full bars are for contralateral leg (c, control) and hatched bars are for hypertrophied leg (h, overload surgery). A, Immunohistochemistry of EDL muscle sections of Ctrl and *Piezo1^ΔEC^* mice for nuclei (DAPI, blue). Scale bars, 100 μm. B, Nuclei density; C, Nuclei-to-fibre ratio. Data are for N = 4-5 mice per group (mean ± S.D.). Superimposed dots are the individual underlying data values for each individual mouse. \**P*<0.05, \*\**P*<0.01, \*\*\**P*<0.001 vs. contralateral leg. Statistical significance was evaluated using 2-way ANOVA followed by Tukey’s HSD post hoc test for multiple comparisons.

**Figure 6:**
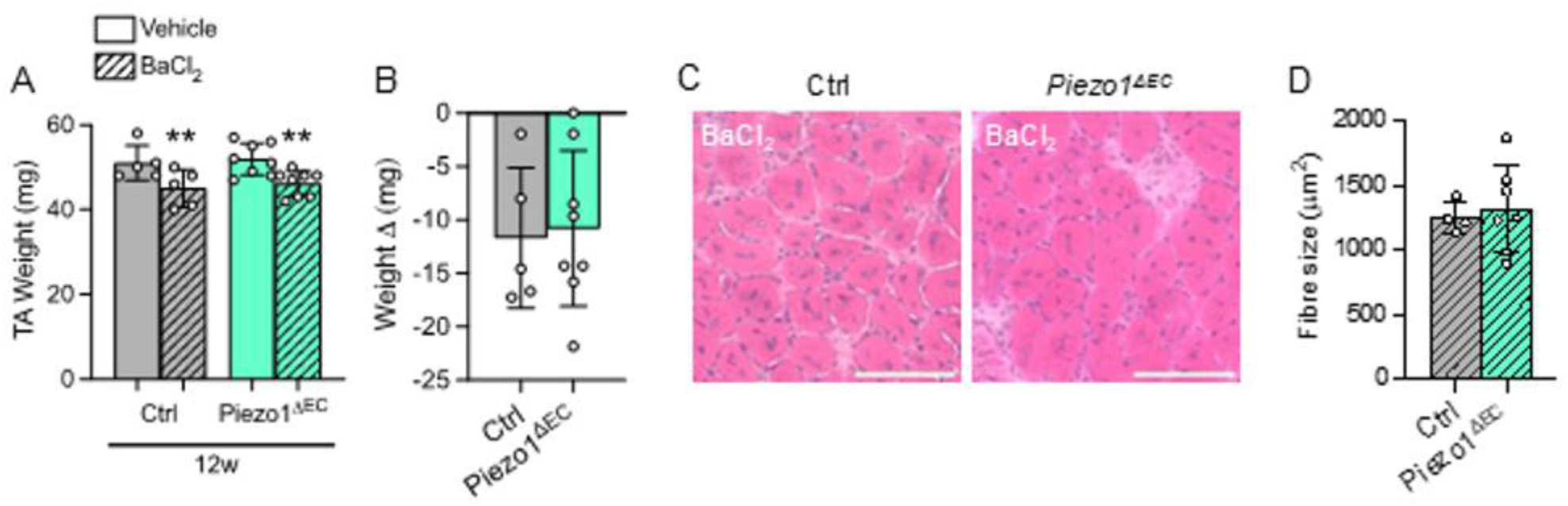
Two-week deletion of endothelial Piezo1 does not change muscle regeneration. Data in grey are for 12-weeks old Ctrl mice and data in colour are for matched *Piezo1^ΔEC^* mice. Full bars are for contralateral leg (vehicle-injected leg) and hatched bars are for barium chloride (BaCl_2_)-injected leg (A-D). A, Tibialis anterior (TA) muscle wet-mass; B, Delta of muscle loss between BaCl_2_-injected leg and vehicle-injected leg. C, Histological evaluation of BaCl_2_-injected TA muscles using H&E staining. Scale bar, 100 μm. D, Fibre size determination using the mean cross-sectional area. Data are for N = 5-8 (A-D) mice per group (mean ± S.D.). Superimposed dots are the individual underlying data values for each individual mouse. \*\**P*<0.01 vs. vehicle injected leg. Statistical significance was evaluated using 2-way ANOVA followed by Tukey’s HSD post hoc test for multiple comparisons.

**Table 1:** Morphometric parameters from Ctrl and Piezo1^ΔEC^ mice submitted to BaCl_2_-induced muscle regeneration. \**P*<0.05, \*\**P*<0.01, \*\*\**P*<0.001 vs. control leg (vehicle-injected leg). Statistical significance was evaluated using 2-way ANOVA followed by Tukey’s HSD post hoc test for multiple comparisons.

| *BaCl<sub>2</sub> injection* | Ctrl<br>(N=5) |  | *Piezo1*<sup>ΔEC</sup><br>(N=8) |  |
| --- | --- | --- | --- | --- |
|  | Vehicle | BaCl<sub>2</sub> | Vehicle | BaCl<sub>2</sub> |
| Body weight, g | 28.31 ± 1.24 |  | 28.19 ± 1.73 |  |
| Tibia length, mm | 18.33 ± 0.57 |  | 18.16 ± 0.47 |  |
| TA weight on body weight ratio | 1.80 ± 0.08 | 1.59 ± 0.13 ** | 1.84 ± 0.11 | 1.64 ± 0.15 ** |
| TA weight on tibia length ratio | 2.78 ± 0.16 | 2.45 ± 0.23 ** | 2.86 ± 0.26 | 2.54 ± 0.18 ** |

### Endothelial Piezo1 may act by regulating total AKT

To explore alternative mechanisms for the reduced load-induced muscle hypertrophy, we applied RNA sequencing to the EDL from Ctrl and *Piezo1*^ΔEC^ mice. Unbiased RNAseq analysis identified 8506 differentially expressed genes (DEGs) for pairwise comparison between the contralateral and hypertrophied EDL in control mice, and 7680 DEGs in *Piezo1*^ΔEC^ mice (Figure S2A). Enriched pathway analysis using KEGG database highlighted common pathways between genotypes such as chemokine signalling pathways (mmu04062), cytokine-cytokine receptor interaction (mmu04060) and extracellular matrix receptor interaction (mmu04512) (Figure S2B). Specifically, results suggested differences in gene expression relating to apoptosis and mitochondrial regulation, stress and inflammatory signalling, and protein homeostasis and growth regulation (Figure S3). We therefore explored changes in protein homeostasis because of prior evidence of its importance in muscle hypertrophy^2^.

A key mechanism to increase protein synthesis and subsequent muscle hypertrophy is increased AKT-mTORC1 signalling^3^. We found no clear differences between groups in the phosphorylation status of mTORC1 or S6 after 2 weeks of overload, and similar degrees of activation in this pathway across genotypes (Table S1). This may be explained by measuring changes at an extended time point of 2 weeks, given increased mTORC1 signalling can be acute and transient following mechanical/exercise stimuli that diminishes over time^37^. However, while there was no change in AKT phosphorylation at its key regulatory residue serine 473^38^ (Figure 7A-C), total AKT after overload increased less in *Piezo1^ΔEC^* mice (Figure 7D). There were no differences between groups in the phosphorylation status of mTORC1 or its downstream target ribosomal protein S6 after overload, with similar degrees of activation in this pathway across genotypes (Table S1). These data suggest that endothelial Piezo1 may act specifically by regulating the abundance of total AKT (protein kinase B), a key kinase regulator of muscle protein synthesis.

**Figure 7:**
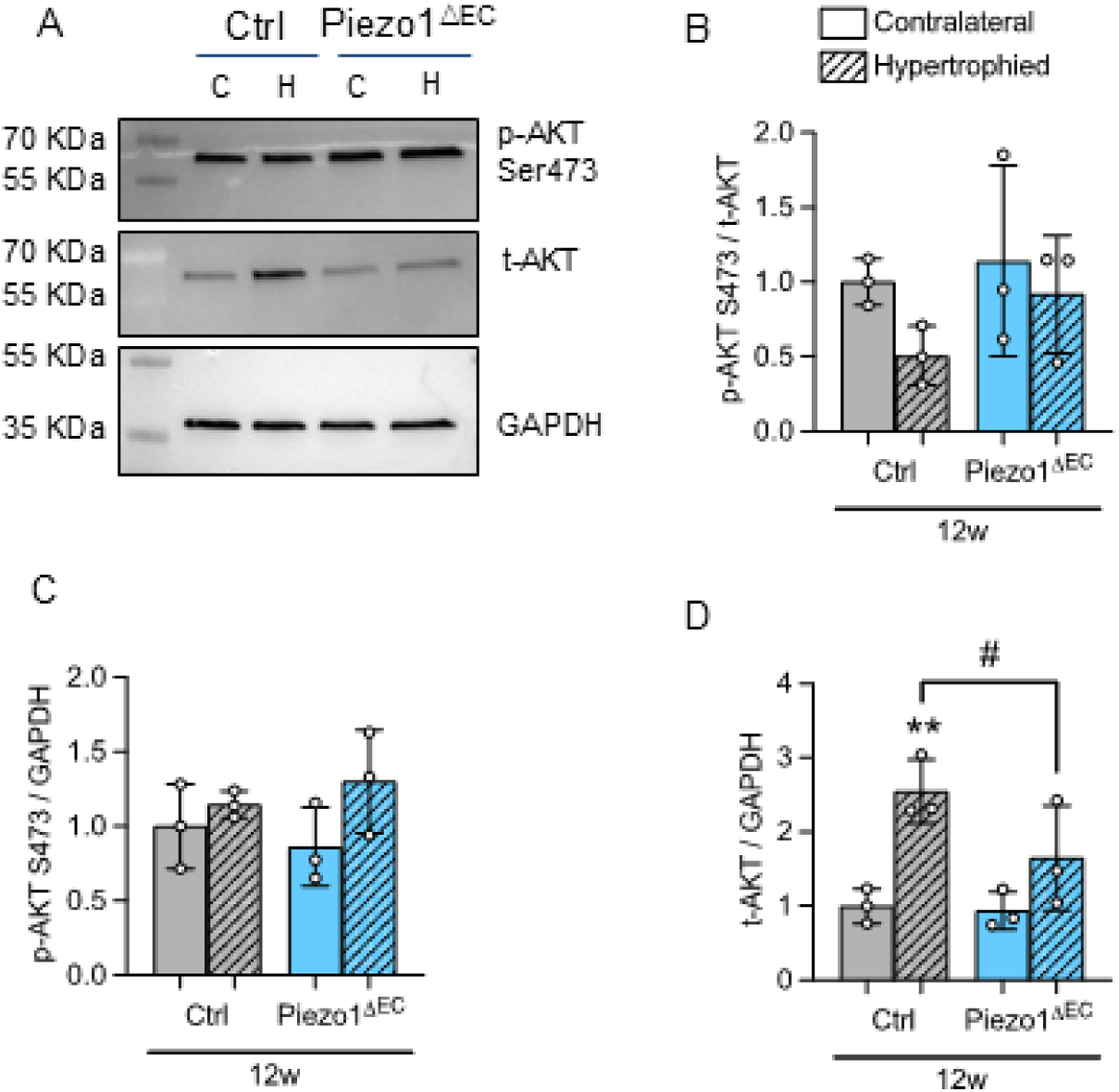
Two-week deletion of endothelial Piezo1 influences mechanical-induced increase in AKT expression. Data in grey are for 12-weeks old Ctrl mice and data in colour are for matched *Piezo1^ΔEC^*mice. For data A-D, full bars are for contralateral leg (c, control) and hatched bars are for hypertrophied leg (h, overload surgery). A, Representative Western blot for AKT phosphorylation at serine 473 (p-AKT) and total AKT (t-AKT) protein in EDL muscle. For data of the type shown in A, quantification of p-AKT relative to t-AKT (B), quantification of p-AKT relative to the housekeeper protein GAPDH (C), quantification of t-AKT relative to the housekeeper protein GAPDH (D). Data are for N = 3 mice per group (mean ± S.D.). Superimposed dots are the individual underlying data values for each individual mouse. \*\**P*<0.01 vs. contralateral leg. ^#^*P*<0.05 vs. ctrl mice. Statistical significance was evaluated using Student’s *t* test (F-H) or 2-way ANOVA followed by Tukey’s HSD post hoc test for multiple comparisons (B-D).

## DISCUSSION

Our results suggest that acute disruption of Piezo1 in the endothelium impairs skeletal muscle growth in response to mechanical loading in mice but is dispensable for muscle regeneration following injury. Endothelial Piezo1 therefore seems instrumental for transducing additional mechanical tension into muscle growth, highlighting a previously unrecognised mechanosensory mechanism in the muscle niche coordinating endothelial-myofibre crosstalk. Furthermore, given that muscle mass is closely matched by enhanced vascular adaptations^3,11,30^, our data highlight this may be achieved through endothelial Piezo1. Although the precise downstream pathways remain to be established, endothelial Piezo1 seems to act in part by regulating the total abundance of AKT protein. Future studies incorporating temporal resolution during the acute post-loading period may help in defining the molecular signals linking this endothelial mechanosensing to skeletal muscle hypertrophy.

The Piezo1 protein channel is a critical biological mechanosensor within the body and involved in many wide-ranging physiological processes. The role of Piezo1 channels regulating skeletal muscle mass and physical performance has gained increasing recognition in recent years, however this seems highly cell-type dependent^39,40^. For example, in mouse models myofiber-specific Piezo1 regulates muscle atrophy in mice via KLF15-IL6 dependent signalling during limb immobilisation^24^, whereas MuSC-specific Piezo1 is necessary for muscle regeneration following localised injury^13,25^. Furthermore, in mice expressing a gain-of-function mutation of Piezo1 in tendons, jumping ability and running speeds are increased and this mutation is more frequent in Jamaican sprinters linking to human performance^22^. Our group have previously reported that endothelial-specific Piezo1 determines physical fitness and is necessary for maintaining long-term muscle capillarity in mice^27^. In this study our data now show a previously unrecognised role for endothelial Piezo1 in regulating muscle growth following mechanical loading.

We found that muscle mass was blunted and myofibre hypertrophy absent in *Piezo1*^ΔEC^ compared to Control mice in response to the well-established functional overload model (synergist ablation)^3^. We therefore propose mechanical loads initiate an increase in Piezo1 signalling within resident muscle endothelial cells (e.g. via increased blood flow and/or stretch), which in turn provides a unique vascular ‘signal’ to optimise normal muscle growth. Although mechanical loads have been linked to baseline muscle capillarity and increased angiogenesis^29–32^, with long-term endothelial Piezo1 deletion decreasing muscle capillarity^27^, the current study suggests an additional early-onset and capillary density-independent mechanosensory mechanism. For example, we observed no differences between Piezo1^ΔEC^ and Control mice in baseline muscle capillarity, although we did find endothelial Piezo1 deletion mildly blunted the angiogenic response (C:F ratio) 2 weeks following mechanical overload. An endothelial mechanosensory adaptation such as this may have evolved to optimise the coupling between muscle growth/mass and blood flow^11^, reducing the risk of adaptation that cannot be supported by sufficient blood supply. In the short term, this endothelial-myofiber coupling may serve to fine tune and maximise muscle growth early in response to mechanical loads, by overcoming potential limitations in cues derived from the local muscle niche and myofiber. However, in the longer term any increase in muscle mass would need to be matched by increases in muscle vascularity (e.g. angiogenesis, blood flow). Interestingly, this may serve as a positive feedback loop such that improved vascular adaptations may then further facilitate acute endothelial-myofiber crosstalk in response to mechanical loading to once again support muscle growth in the short term.

There is relatively poor understanding of how an endothelial-myofibre partnership could promote muscle growth induced by mechanical loads^12^. Our data exclude MuSC-dependent myonuclear accretion as a primary mechanism^32–34^, as we found similar increases after overload between *Piezo1*^ΔEC^ and Ctrl groups. No differences were found between groups in muscle regenerative capacity following acute injury, a process dependent upon MuSCs^36^. Instead, we propose that the impaired hypertrophic response observed in *Piezo1*^ΔEC^ mice may be attributable, at least in part, to attenuated myofibre protein synthesis following mechanical loading^3^ and this may be due to an effect on total AKT. We speculate that in response to mechanical loads in skeletal muscle, endothelial Piezo1 may provide an early signal to increase mTORC1 signalling to enhance rates of protein synthesis, which in the long-term may also influence AKT regulation to collectively influence muscle growth. Although we did not observe clear differences between groups in terms of phosphorylated AKT-mTORC1 signalling at 14 d, this could reflect the transient nature of these signalling events, which typically occur during the early stages of adaptation and may not be captured at the later time points we used^37^. Such effects may have been further obscured by our more physiologically relevant but modest EDL growth model compared to the more severe plantaris overload approach^41^. Future studies examining earlier post-loading time points, together with direct assessments of muscle protein synthesis using approaches such as puromycin incorporation, may help determine whether endothelial Piezo1 regulates the acute anabolic response to mechanical loading.

Taken together, our findings identify a previously unrecognised mechanism in endothelial cells that contributes towards load-induced muscle growth. Further work is required to increase knowledge on the endothelial Piezo1 role in skeletal muscle homeostasis, determine downstream signalling pathways, and explore whether this mechanism could be a target for treating low muscle growth and atrophy in disease and ageing.

## Supplementary Figures and Tables

**Supplementary Figure 1:**
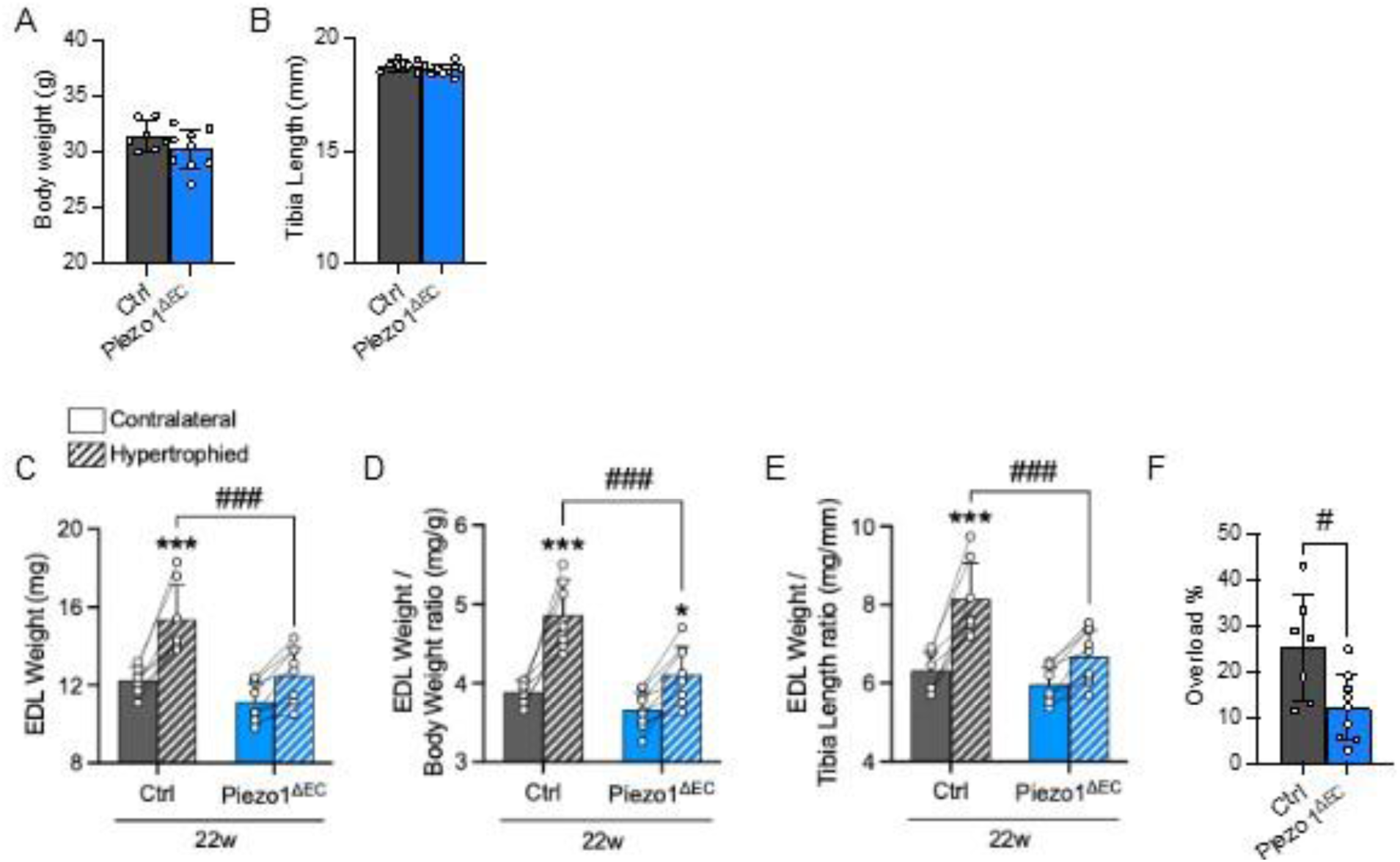
Ten-week deletion of endothelial Piezo1 blunts overload-induced myofibre hypertrophy. Data in dark grey are for 22-weeks old Ctrl mice and data in dark blue are for matched Piezo1^ΔEC^ mice. Plain colour is for contralateral leg (sham surgery) and stripped colour is for hypertrophied leg (overload surgery). A, Body weight; B, Tibia length; C, Dry EDL muscle weight; D, EDL weight to body weight ratio; E, EDL weight to tibia length ratio; F, Percentage of overload degree of hypertrophied leg vs. contralateral leg. Data are for N = 7-9 mice per group (mean ± S.D.). Superimposed dots are the individual underlying data values and lines are direct comparison between contralateral and hypertrophied legs for each individual mouse. \**P*<0.05, \*\*\**P*<0.001 vs. contralateral leg, ^#^*P*<0.05, ^###^*P*<0.001 vs. ctrl mice. Statistical significance was evaluated using Student’s *t* test (A-B, F) or 2-way ANOVA followed by Tukey’s HSD post hoc test for multiple comparisons (C-E).

**Supplementary Figure 2:**
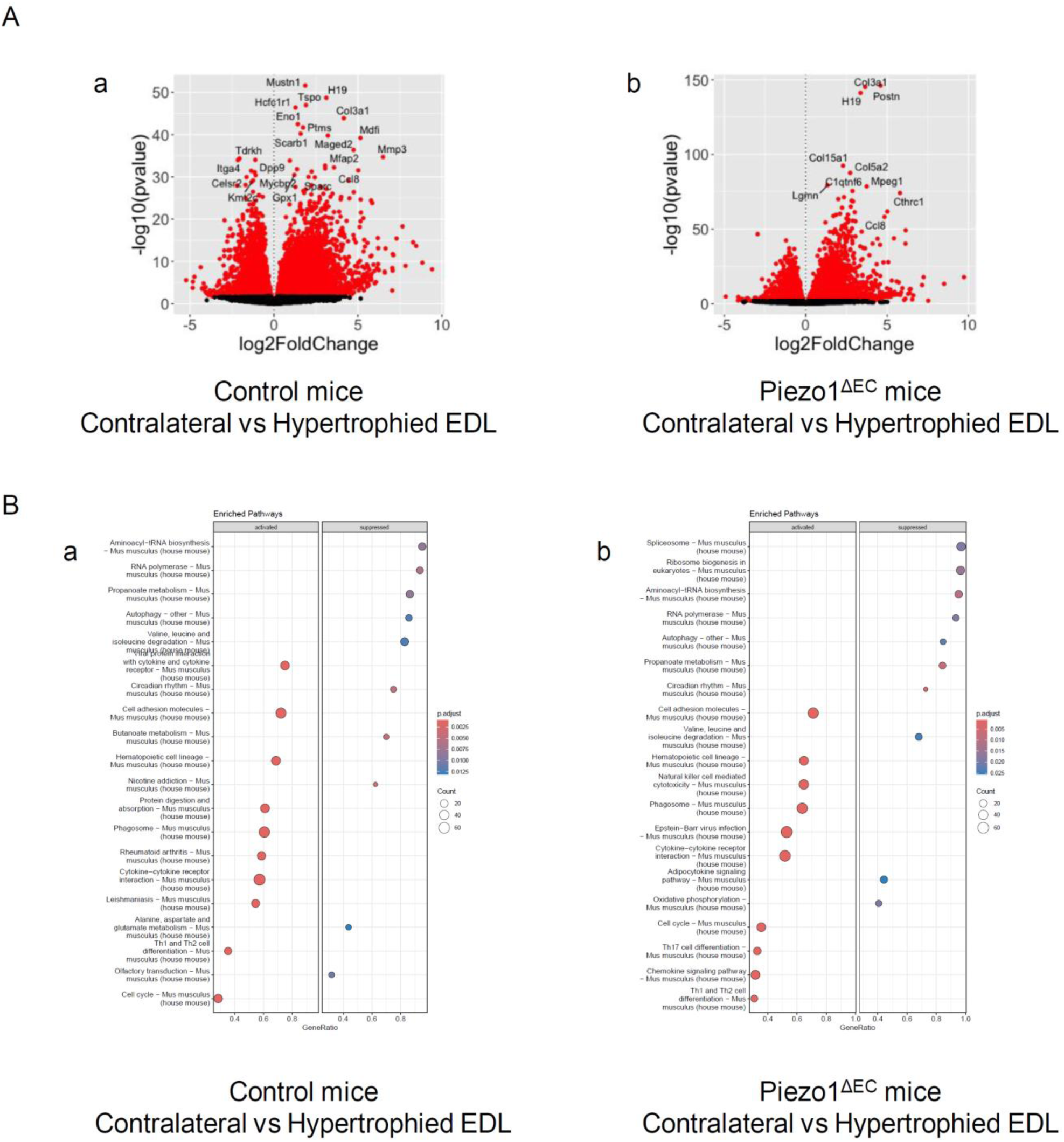
RNAseq data analysis. A, Differentially expressed genes (DEGs); differential expression analysis was performed using DESeq2. A DEG is determined by adjusted P-value<0.05 which corrected using Benjamini-Hochberg method. Pairwise comparisons were conducted such as (a) Control mice – Contralateral vs Hypertrophied side (8506 DEGs identified) and (b) Piezo1^ΔEC^ mice – Contralateral vs Hypertrophied side (7680 DEGs identified). B, Enriched pathway analysis using KEGG database. Pathways that achieved adjusted P-value<0.05 were considered significant. Top 20 enriched pathways presented for (a) Control mice – Contralateral vs Hypertrophied side and (b) Piezo1^ΔEC^ mice – Contralateral vs Hypertrophied side.

**Supplementary Figure 3:**
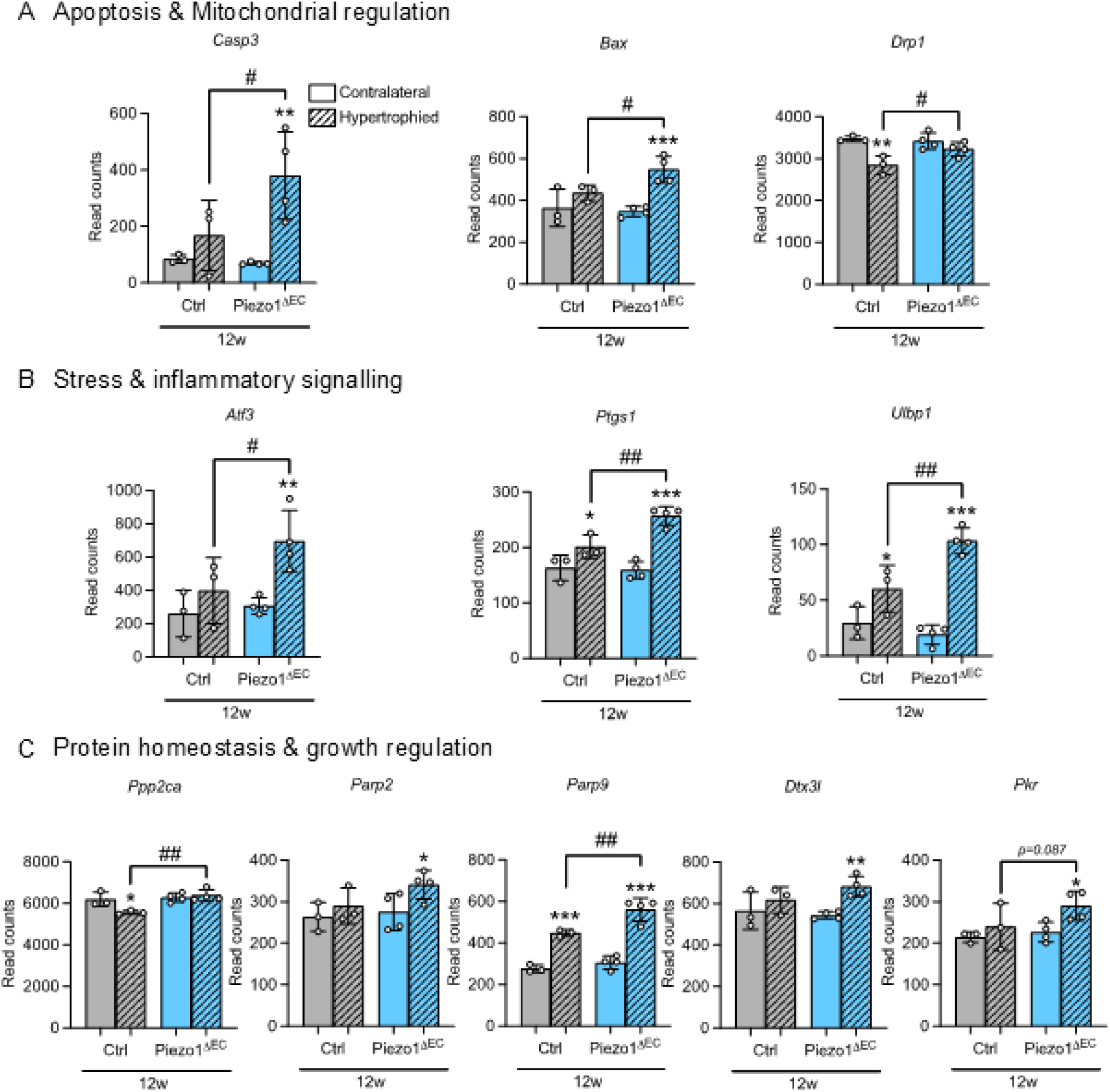
Gene expression from Ctrl and Piezo1^ΔEC^ mice submitted to muscle overload. The genes indicated were selected as markers of apoptosis and mitochondrial regulation, stress and inflammatory signalling, protein homeostasis and growth regulation. Gene expression was determined by bulk RNA sequencing of EDL muscle from Ctrl and Piezo1^ΔEC^ mice for both contralateral and hypertrophied leg. The abundance of mRNA is presented as read counts. All displayed values are mean ± S.D. Statistical significance was evaluated using 2-way ANOVA followed by Tukey’s HSD post hoc test for multiple comparisons.

**Table S1:** Western blot quantification of protein synthesis pathway from Ctrl and Piezo1^ΔEC^ mice submitted to muscle overload. \**P*<0.05 vs. contralateral leg. Statistical significance was evaluated using 2-way ANOVA followed by Tukey’s HSD post hoc test for multiple comparisons.

|  | Ctrl (N=3) |  | Piezo1<sup>ΔEC</sup> (N=3) |  |
| --- | --- | --- | --- | --- |
|  | C | H | C | H |
| p-mTOR / t-mTOR | 1.00 ± 0.04 | 0.88 ± 0.07 | 0.97 ± 0.12 | 1.03 ± 0.35 |
| p-mTOR / GAPDH | 1.00 ± 0.14 | 1.26 ± 0.38 | 0.34 ± 0.34 | 1.45 ± 0.47 |
| t-mTOR / GAPDH | 1.00 ± 0.09 | 1.47 ± 0.36 * | 0.96 ± 0.27 | 1.40 ± 0.06 * |
| p-S6 / t-S6 | 1.00 ± 0.38 | 1.79 ± 1.41 | 1.38 ± 0.61 | 1.54 ± 1.00 |
| p-S6 / GAPDH | 1.00 ± 0.27 | 4.08 ± 4.14 | 1.45 ± 0.86 | 2.87 ± 1.18 |
| t-S6 / GAPDH | 1.00 ± 0.13 | 1.95 ± 0.57 * | 0.95 ± 0.26 | 1.98 ± 0.51 * |

**Table S2:** Antibodies for immunoblotting.

| Category | Protein / Target | Host | Dilution | Catalogue reference | Source |
| --- | --- | --- | --- | --- | --- |
| Primary antibody | p-Akt (Ser473) | Rabbit | 1:1000 | #4060 | Cell Signaling |
| Primary antibody | t-Akt | Rabbit | 1:1000 | #9272 | Cell Signaling |
| Primary antibody | p-mTOR (Ser2448) | Rabbit | 1:1000 | #5536 | Cell Signaling |
| Primary antibody | t-mTOR | Rabbit | 1:1000 | #2983 | Cell Signaling |
| Primary antibody | p-S6 (Ser240/244) | Rabbit | 1:1000 | #2215 | Cell Signaling |
| Primary antibody | t-S6 | Rabbit | 1:1000 | #2217 | Cell Signaling |
| Primary antibody | GADPH HRP | Mouse | 1:10000 | ab105428 | Abcam |
| Secondary antibody | Rabbit | Donkey | 1:5000 | 711-036-152 | Jackson Immunoresearch |

## Funding

TSB was supported by a BBSRC International Partnering Award.

## Disclosures

None

## Notes

### Competing Interest Statement

The authors have declared no competing interest.

### Summary of Updates

Addition of Figure 7 and Supplementary Figures 2 and 3, to add information on the underlying molecular mechanisms. Overall interpretation updated.

